# A genome catalogue of lake bacteria across watershed land use gradients at a continental scale

**DOI:** 10.1101/2022.08.12.503676

**Authors:** Rebecca E. Garner, Susanne A. Kraemer, Vera E. Onana, Maxime Fradette, Marie-Pierre Varin, Yannick Huot, David A. Walsh

## Abstract

Lakes are heterogenous ecosystems inhabited by a rich microbiome whose genomic diversity is poorly defined. We present a continental-scale study of metagenomes representing 6.5 million km^2^ of the most lake-rich landscape on Earth. Analysis of 308 Canadian lakes resulted in a metagenome-assembled genome (MAG) catalogue of 1,008 mostly novel bacterial genomospecies. Lake trophic state was a leading driver of taxonomic and functional diversity among MAG assemblages, reflecting the responses of communities profiled by 16S rRNA amplicons and gene-centric metagenomics. Coupling the MAG catalogue with watershed geomatics revealed terrestrial influences of soils and land use on assemblages. Agriculture and human population density were drivers of turnover, indicating detectable anthropogenic imprints on lake bacteria at the continental scale. The sensitivity of bacterial assemblages to human impact reinforces lakes as sentinels of environmental change. Overall, the LakePulse MAG catalogue greatly expands the freshwater genomic landscape, advancing an integrative view of diversity across Earth’s microbiomes.

## Main

Freshwater bacteria are a diverse component of lake ecosystems and are central to biogeochemical cycles and water quality^1–3^. Collectively, Earth’s millions of lakes exhibit immense environmental heterogeneity, reflecting a range of influences from regional variations in geology, landform, and climate to local compositions of catchment runoff^4^. Lakes integrate terrestrial carbon and other materials from the catchment and airshed, to an extent that lakes are considered an axis of the global carbon cycle^5^ and sentinels of environmental change^6^. The growing number of metagenomic studies is providing important insights into the ecology and evolution of freshwater bacteria in the context of changing environmental conditions^7^. However, most studies are focused on a limited number of lakes, restricting our understanding on how environmental variation influences the structure and function of bacterial assemblages across lake ecosystems. Elucidating freshwater bacterial genomic diversity is critical given the myriad anthropogenic pressures on lakes. Lake warming^8^, eutrophication^9^, deoxygenation^10^, salinization^11^, chemical contamination, and other emergent stressors^12^ are expected to modify the freshwater microbiome, but in ways that we are only beginning to understand^13^.

Genome-resolved metagenomics enables the reconstruction of composite genomes from microbial populations^14^. MAGs supply reference genomes for uncultivated taxa, revealing enormous microbial dark matter^15^. Currently, MAG analysis is a standard approach to link functions to populations in complex communities. Lake studies applying genome-resolved metagenomics have brought insights into the evolution^16, 17^, metabolism^18–20^, and succession^21, 22^ of freshwater bacteria. Recently, the approach has been applied at larger scale to generate genomic catalogues from marine^23^, terrestrial^24^, and host-associated microbiomes^25–27^, culminating in a genome catalogue of Earth’s microbiomes (GEM)^28^. Although impressive in its collection of habitats, GEM encompasses very few freshwater lakes (<30), compared to the immense abundance and heterogeneity worldwide. Further efforts have begun to elucidate lake genome diversity in stratified water columns^29^. An essential next step is to expand the representation of freshwater lakes in the global genomic catalogue.

Here, we introduce a genome-resolved perspective of freshwater bacterial diversity across 6.5-million km^2^ of Canada, the most lake-rich landscape on Earth^30^. This study is a component of the NSERC Canadian LakePulse Network^31^, a large-scale coordinated effort to assess the health status of lakes using standardized sampling and analysis methods with a major focus on microbial communities^32–40^. Through metagenome co-assembly and binning, we reconstructed 1,184 high-and medium-quality MAGs from 308 lakes reflecting expansive gradients in limnological and watershed conditions. The LakePulse MAG catalogue was then used to investigate the structure and function of lake bacterial assemblages at continental scale. In addition to clarifying the functional repertoires of freshwater taxa, we pursued a novel analysis of the metabolic variation among MAG assemblages across lake physicochemical and other environmental gradients. The consideration of geomatics variables in the LakePulse survey further allowed us to report on watershed factors influencing bacterial assemblages across the land-water interface, including detectable land use effects signaling evidence of human impact. In resolving genomic diversity from the largest representation of lakes across a temperate to subarctic range, the LakePulse MAG catalogue forms a key resource for investigating the freshwater microbiome on a rapidly changing planet.

## Results

### The LakePulse MAG catalogue

To produce the MAG catalogue, we generated metagenomes from the surface waters of 308 lakes in 12 Canadian ecozones (**Fig. 1**; **Fig. S1**; **Table S1**). Lakes represented a wide range of physicochemical conditions, productivity, morphometry, climate, and land use within watersheds^41^ (**Fig. S2)**. At the continental scale (43 – 68 °N, 62 – 141 °W), a full range of ultraoligotrophic to hypereutrophic lakes were present in the survey (**Fig. 1A**). Comparison of lake features by principal component analysis (PCA) revealed large-scale spatial patterns in lake and watershed characteristics (**Fig. 1B**). Western Canadian lakes were the deepest and most oligotrophic on average, and often set in watersheds with high proportions of natural landscapes and harvested forests. Northern Canadian lakes experienced the coldest climates and lowest land use pressures. Central Canada was comprised of the agriculturally intensive Prairies and Boreal Plains ecozones; compared to other Canadian regions, central Canadian lakes were generally shallow, alkaline, nutrient-and ion-rich, and highly productive. Eastern Canadian lakes had the warmest surface waters and on average the most extensive built environments in their watersheds. Agriculture was also a common land use in eastern Canada, particularly within the Mixedwood Plains ecozone. Lakes in watersheds with the largest population centres were in the Vancouver and Toronto metropolitan areas within the Pacific Maritime and Mixedwood Plains ecozones, respectively.

**Figure 1.**
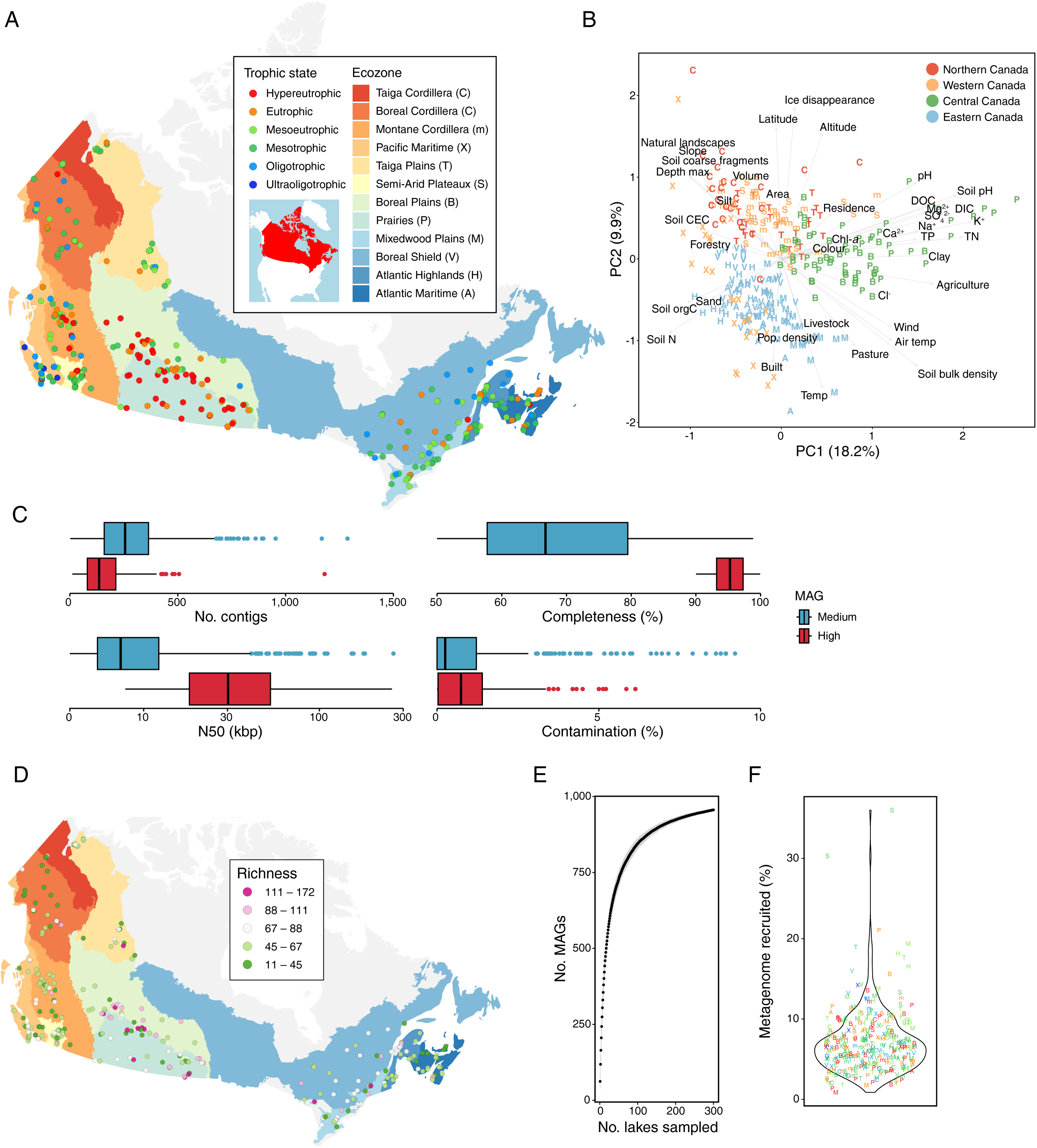
The LakePulse MAG catalogue: geographical and environmental scope, quality, and patterns of local-scale diversity. **(A)** Map of 308 sampled lakes from across a broad trophic state gradient (shown as point colours) in 12 Canadian ecozones (shown as base map colours). **(B)** Principal component analysis illustrating the variation in environmental features among sampled lakes. The letter symbols (see panel A for legend) and colours of each point represent ecozones and geographic regions of Canada, respectively. Arrows show the contributions of environmental variables to PC loadings, except for latitude which was passively plotted. **(C)** Distributions of MAG characteristics for medium-and high-quality MAGs. **(D)** MAG richness within lakes. Point colours represent richness, which is the number of MAGs detected at each site based on positive TAD_80_ values (see panel A for ecozone colour legend). **(E)** MAG accumulation curve (i.e. the number of new genomes detected) in a random ordering of lakes. **(F)** Fraction of LakePulse metagenomes captured within MAGs (measured as the percentage of each metagenome recruited to the MAGs). Point colours represent lake trophic states and letters represent ecozones (see legend of panel A).

We aimed to capture a broad phylogenetic and phenotypic diversity of microorganisms (e.g., organism size and lifestyle) in the LakePulse survey. To this end, lake metagenomes were generated from biomass collected across a wide size range (0.22 – 100 μm). Eleven metagenome co-assemblies representing the 12 ecozones were generated (Boreal and Taiga Cordilleras lakes were combined). Though computationally intensive to produce, co-assemblies facilitated differential coverage binning, ensuring higher-quality MAGs^42^. In total, contig binning of each co-assembly recovered 1,184 MAGs exceeding MIMAG medium-quality standards^43^ through an additional <10% strain heterogeneity criterion (**Table S2**). Between 28 – 233 MAGs were generated for each ecozone (**Fig. S3**). MAGs were dereplicated across ecozones (ANI ≥95%), resulting in a genomospecies-level set of 1,008 MAGs. MAGs were categorized as 136 high-and 872 medium-quality drafts based on MIMAG standards^43^, excluding rRNA gene requirements (**Fig. 1C**). MAGs ranged widely in size (0.35 – 10.39 Mbp), GC content (23.1 – 75.9%), and coding density (78.4 – 96.8%) (**Table S2**). Single metagenome assemblies produced only 563 bins passing our quality control (**Table S1**); these represented just 202 lakes and 476 genomospecies, over one-third of which were already present in the catalogue. This comparison validated our more intensive co-assembly approach to reconstructing lake bacterial genomic diversity.

To map MAG biogeography across lakes, we calculated the central 80% truncated average depths of coverage (TAD80) from fragment recruitment^21^. Based on TAD80, observed richness within lakes varied between 11 – 172 MAGs, with diversity hotspots tending to occur in the shallow, nutrient-rich lakes of the Boreal Plains and Prairies ecozones of central Canada (**Fig. 1D**). A species accumulation curve showed that hundreds of lakes were required to adequately sample genomes across the heterogenous Canadian landscape, highlighting the importance of analyzing lake metagenomes across large geographic and environmental gradients to capture the breadth of local lake bacterial diversity and ecosystem function (**Fig. 1E**). MAGs captured on average 7.1% (range 0.9 – 36.0%) of metagenome reads per lake (**Fig. 1F**), signifying that the MAG catalogue is a reasonable but incomplete representation of bacterial genome diversity in the lakes. Currently, the LakePulse MAG catalogue represents a significant expansion in the availability of freshwater genomes and range of lake systems to date.

### Phylogenetic and metabolic diversity of novel lake bacterial species

The LakePulse MAG catalogue represented a phylogenetically rich set of genomes (**Fig. 2A**; **Table S2**) exhibiting high taxonomic novelty. Overall, 96% of MAGs were novel candidate species and over a third (349) belonged to novel genera (**Fig. 2B**), substantially expanding the known genomic diversity of lake bacteria. The largest collection of MAGs was recovered from *Bacteroidota*, a group that plays a major role in organic particle degradation in marine settings^44^ but is much less studied in freshwater systems^3^. A third of *Bacteroidota* MAGs (80 of 275) represented new genera (**Table S3**), presenting a rich genomic resource for future targeted research on freshwater *Bacteroidota* ecology and metabolism, including potential zooplankton associations^45^. *Actinobacteriota* and *Gammaproteobacteria* were the next most MAG-rich groups and included lineages of common and abundant freshwater bacteria (e.g., *Rhodoluna*, *Polynucleobacter*, and *Limnohabitans*). Other freshwater taxa that were well represented in the MAG catalogue included *Verrucomicrobiota*, *Planctomycetota*, *Patescibacteria*, and *Myxococcota*. Intriguingly, we identified 38 MAGs from *Bdellovibrionota*, almost all of which represented new genera (**Table S3**). *Bdellovibrionota* are described as obligate predators of Gram-negative bacteria^46^ and their role in freshwater food webs warrants further attention.

**Figure 2.**
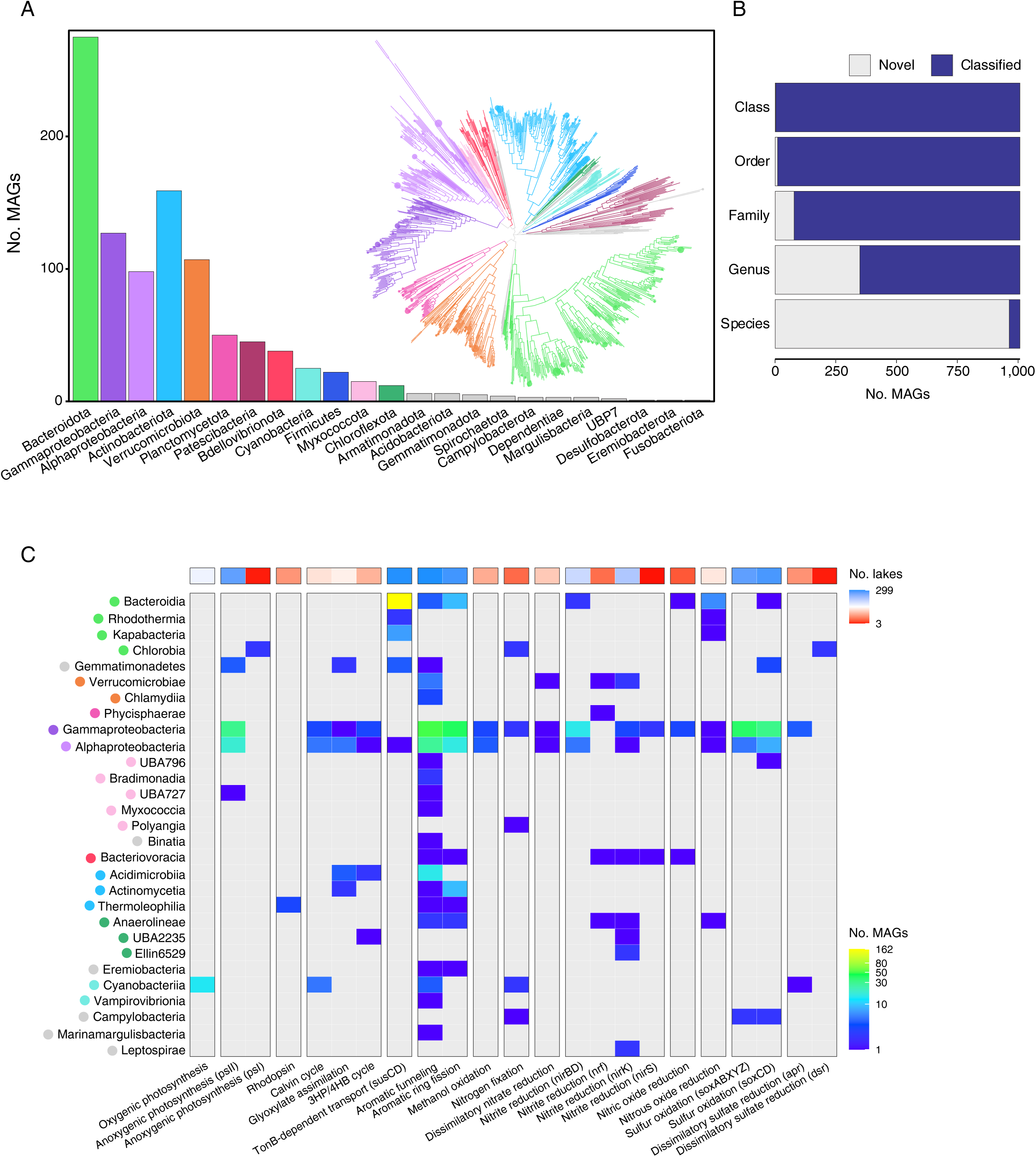
Taxonomic, phylogenetic, and functional diversity of the LakePulse MAG catalogue. **(A)** Number of MAGs assigned to each phylum and phylogenetic tree of MAGs constructed from 120 bacterial marker genes. **(B)** Taxonomic novelty of the LakePulse MAG catalogue. Bar plots are divided by colour to show the number of novel (i.e. unclassified) and classified MAGs from class to species level. **(C)** Functional diversity across taxonomic orders. The lower heat map shows the number of MAGs within each class containing genetic markers for a selection of metabolic functions (grey = no evidence; point colours next to class names indicate the phylum-level affiliation of MAGs shown in panel A). The upper heat map shows the number of lakes containing MAGs with evidence of metabolic functions.

To assess whether the MAGs captured a representative phylogenetic diversity of lake bacteria, we compared taxonomy between MAGs and 16S rRNA gene amplicon sequence variants (ASVs). MAGs and ASVs overlapped in their 10 most diverse phyla, though ASVs exhibited an expected long tail of rare diversity (**Fig. S4A**). Moreover, the three most MAG-rich orders (*Chitinophagales*, *Burkholderiales*, and *Flavobacteriales*) were in the top four among ASVs (**Fig. S4B**). The largest differences in diversity across datasets were within *Bacteroidota* and *Actinobacteriota*, which were slightly more represented as a percentage of MAGs. In contrast, *Patescibacteria* exhibited higher ASV diversity and were potentially underrepresented in the MAGs because of low or divergent marker gene content^47^. Overall, the LakePulse MAG catalogue encompasses broad phylogenetic diversity reflective of 16S rRNA genes, making it a useful resource for further taxon-focused research.

Bacteria contribute to lake ecosystem function through a diversity of energy-capturing and carbon cycling metabolisms that regulate biogeochemical cycles. To provide a broad overview of MAG metabolic diversity, we explored the distributions of select metabolic modules across the MAG catalogue (**Fig. 2C**; **Table S2**). Oxygenic photosynthetic *Cyanobacteria* and anoxygenic photosynthetic *Chlorobia* were present, while bacteriorhodopsins were identified in *Thermoleophilia*. Photosystem II (PS-II) was the most widespread phototrophic module in the MAGs, including in numerous *Alphaproteobacteria*, *Gammaproteobacteria*, and *Gemmatimonadota* MAGs. In combination, PS-II-encoding MAGs were identified in 95% of lakes, indicating a diverse bacterial contribution to freshwater phototrophy. We identified three of the six known autotrophic carbon fixation pathways^48^, including the Calvin-Benson-Bassham (CBB) cycle, the 3-hydroxypropionate/4-hydroxybutyrate (3HP/4HB) cycle, and the glyoxylate assimilation pathway. The CBB cycle was linked to *Proteobacteria* and *Cyanobacteria* MAGs present in 127 lakes (42%), demonstrating that non-algal carbon fixation is also common in lakes.

Heterotrophic degradation of complex algal and terrestrial organic matter is a central ecological function of lake bacteria. An abundance of *Bacteroidota* and *Gemmatimonadota* MAGs were equipped with SusCD-mediated glycan transport systems (**Fig. 2C**; **Table S2**), suggesting broad potential for polysaccharide transport and breakdown. Aromatic compound degradation pathways associated with the metabolism of plant-derived lignin were common within *Alphaproteobacteria* and *Gammaproteobacteria*, but also broadly distributed across the MAG phylogeny (**Fig. 2C**; **Table S2**). In addition to heterotrophy, many *Burkholderiales* MAGs appeared to augment their energy metabolism through Sox-mediated sulfur oxidation pathways. We did not find evidence for methane or ammonia oxidation within the MAG catalogue. However, energy conservation via methanol oxidation and redox cycling of nitrogen oxides was evident. Overall, these findings highlight that the MAG catalogue captures a broad diversity of metabolisms for in-depth comparative analysis on functional contributions to lake ecosystems.

### Lake conditions shape the structure and function of bacterial assemblages

Biogeographical mapping across Canadian lakes revealed that MAGs occupied on average 20 lakes (7%) and as many as 190 lakes (63%) (**Fig. 3A**). Thus, the co-assembly-resolved MAG catalogue appeared to comprise relatively widely distributed bacteria. *Patescibacteria* MAGs were among those with the most restricted ranges, suggesting relatively specialist lifestyles, perhaps dependent on host ranges^49^ or input from groundwater environments where they thrive^50^. Notable biogeographic patterns were observed when MAGs were aggregated at the class to phylum levels (**Fig. S5**; **Fig. S6**). Phyla previously determined to be common in lakes (e.g., *Bacteroidota*, *Proteobacteria*, and *Actinobacteriota*) were broadly distributed across the continent. Other groups were relatively restricted to distinct regions. For example, *Bacilli* and *Campylobacteria* MAGs were more common in Prairies lakes than elsewhere, perhaps in response to high nutrients or extensive agriculture within watersheds (**Fig. S5**; **Fig. S6**).

**Figure 3.**
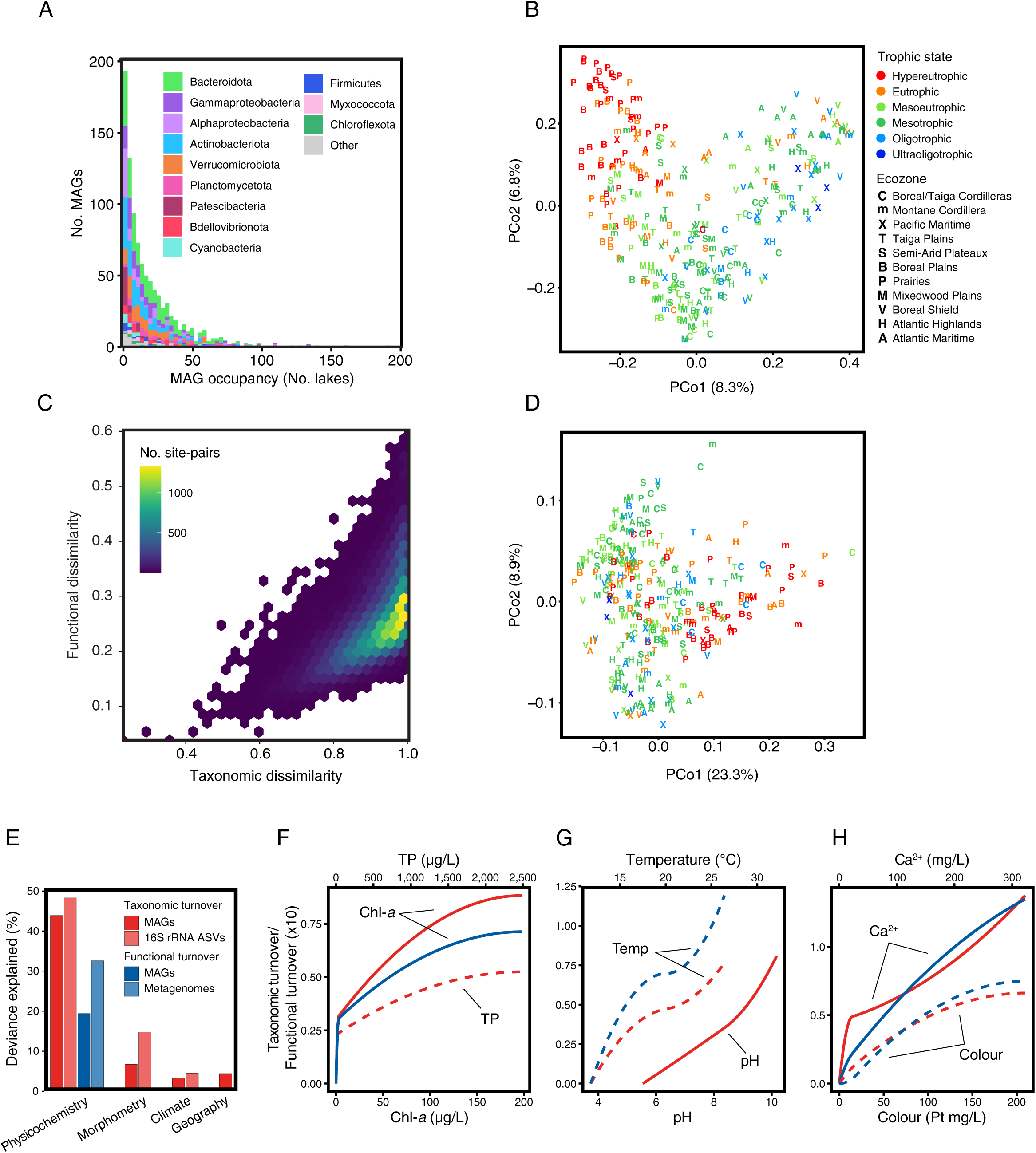
Niche breadth and environmental filtering of MAGs based on taxonomic and functional biogeography. **(A)** Frequencies of MAG incidence (i.e. the number of lakes occupied by each MAG). Colours show MAG phylum-level assignments. **(B)** Principal coordinate analysis (PCoA) showing the variation in MAG taxonomic composition among lakes based on Jaccard dissimilarities. Point colours represent lake trophic state and letter symbols represent ecozones. **(C)** Relationship between functional and taxonomic diversity among MAG assemblages in different lakes. The colour gradient shows the density of pairwise comparisons. **(D)** PCoA showing the variation in functional gene content among MAG assemblages based on Jaccard dissimilarities. Point colours and letter symbols are described in the legend of panel B. **(E)** Percent deviance explained by generalized dissimilarity models (GDMs) fitting the responses of taxonomic or functional MAG assemblages or community profiles to physicochemical, morphometric, geographic, or climatic gradients. (**F-H)** Partial effects of **(F)** total phosphorus (TP) and chlorophyll-*a* (Chl-*a*), **(G)** temperature and pH, and **(H)** calcium and lake colour on the taxonomic and functional turnover of MAG assemblages across lakes. Curve shapes represent the rate of turnover and the maximum height of curves represents the magnitude of turnover across environmental gradients. Line colours represent different MAG assemblage response data (red = taxonomic turnover, blue = functional turnover) and line types (solid or dashed) represent different environmental explanatory variables.

We next used the MAG catalogue to investigate biogeographic patterns of taxonomic and functional diversity across continental-scale environmental gradients. Taxonomic variation among MAG assemblages was correlated with lake trophic state, indexed by total phosphorus (TP) concentration (correlations with principal coordinate analysis [PCoA] axes: *r_1_* = -0.32; *r_2_* = 0.33) (**Fig. 3B**). ASV community taxonomic profiles were likewise distinguished by trophic state (*r_1_* = 0.41) (**Fig. S7A**), demonstrating that MAGs were reasonably representative of bacterial community variation. To assess functional variation, we generated lake functional profiles by aggregating the collection of metabolic KEGG orthologs (KOs) encoded by the MAG assemblage within each lake. Bacterial taxonomic and functional dissimilarities exhibited a positive, but variable relationship (**Fig. 3C**). PCoA of functional dissimilarities revealed a distribution of MAG assemblages along axis 1 related to lake trophic state (*r_1_* = 0.23) (**Fig. 3D**) that reflected the relationship exhibited by metagenome functional profiles (*r_1_* = 0.40; *r_2_* = -0.33; metagenome functional profiles form the basis of a dedicated study [Onana *et al*., in prep.]) (**Fig. S7B**). These results suggest that nutrient availability and productivity play leading roles in structuring the taxonomic and functional composition of lake bacterial assemblages at continental scale.

To further elucidate the factors influencing bacterial diversity across lakes, we evaluated the importance of lake physicochemistry, morphometry, geography, and climate conditions using generalized dissimilarity models (GDMs)^51^. GDMs of membership (Jaccard dissimilarities) explained more deviance compared to structure (Bray-Curtis dissimilarities) (**Table S4**). As membership is more stable within lake assemblages than structure due to the seasonal succession of bacterial populations, it follows that membership is better predicted across large spatial scales. Lake physicochemistry explained the largest amounts of taxonomic (MAGs 43.9%; ASVs 48.3%) and functional (MAGs 19.4%; metagenomes 32.6%) variation among MAG assemblages and community profiles (**Fig. 3E**; **Table S4**). The consistency in deviance explained by taxonomic or functional models showed that MAG assemblages reliably reflected the significant environmental filtering of the broader community. The influence of trophic state was evident as taxonomic and functional turnover across MAG and broader assemblages was consistently responsive to chlorophyll-*a* (Chl-*a*) concentration, a measure of algal biomass (**Fig. 3F**; **Fig. S8A**). MAG taxonomic, but not functional, turnover was significantly related to TP (**Fig. S8A**), suggesting metabolic differences among bacterial assemblages are driven more by the composition of organic matter produced by phytoplankton than by direct nutrient availability. Turnover was most rapid at low Chl-*a* and TP, suggesting rapid shifts in bacterial assemblage composition across oligotrophic to mesotrophic lakes (**Fig. 3F**).

In addition to the effects of trophic state, GDMs provided insights into other physicochemical factors shaping lake bacterial assemblages at continental scale. MAG and community taxonomic turnover was significantly associated with pH, and MAG functional turnover was additionally predicted by lake surface temperature (**Fig. 3G**; **Fig. S8A**). Temperature and pH are often considered master variables driving large-scale spatial patterns in marine^52^ and soil^53^ communities, respectively. As aquatic systems that are heavily influenced by their terrestrial surroundings, it follows that lake bacterioplankton assemblages are influenced by a combination of temperature and pH.

Remarkably, calcium concentration was the strongest predictor of both taxonomic and functional turnover across MAG assemblages and community-level taxonomy (**Fig. 3H**; **Fig. S8A**). A shift in bacterial ecotype diversity along a freshwater calcium gradient was previously reported^54^, but how calcium directly influences assemblage composition is unclear. The effect is perhaps indirectly linked to interactions among plankton, as low calcium was previously shown to limit zooplankton growth^55^, potentially eliciting trophic cascades through the lake food web. Further investigation of the mechanisms and implications of the relationship between calcium and bacterial assemblages is warranted, particularly as lake calcium decline is a legacy effect of emissions-linked acid rain^56^, while calcium levels may be increasing elsewhere through winter road salt application^57^.

The demonstration that lake physicochemistry shapes the functional composition of MAG assemblages prompted a deeper analysis focused on specific metabolic categories. We generated GDMs for different KEGG metabolic categories implicated in energy and nutrient cycling. As with the full functional profiles, lake physicochemistry consistently explained more deviance than morphometry, geography, or climate (**Fig. 4A**). Interestingly, explained deviance varied by metabolic category, and variation within carbohydrate metabolism was the best predicted by physicochemistry (**Fig. 4A**). Significant predictors of carbohydrate metabolism included Chl-*a* concentration and lake colour (**Fig. 4B**; **Fig. S8B**). Lake colour reflects terrestrial organic matter input from the watershed. Hence, variation in carbohydrate metabolism across bacterial assemblages appears to be linked to the quantity and composition of autochthonous (algal) and allochthonous (plant) organic matter in lakes. We note that lipid metabolism was the only additional metabolic category where turnover was predicted by Chl-*a* and colour (**Fig. 4B**; **Fig. S8B**). Subsequent PCoAs of carbohydrate and lipid metabolisms in lakes showed strong relationships with trophic state and large variation across the eutrophic to hypereutrophic spectrum (**Fig. 4C-D**). Overall, these results suggest that a diverse and dynamic organic matter pool characterized by punctuated inputs or resuspensions of nutrients in eutrophic to hypereutrophic systems may be a key driver of metabolic diversity within the lake microbiome.

**Figure 4.**
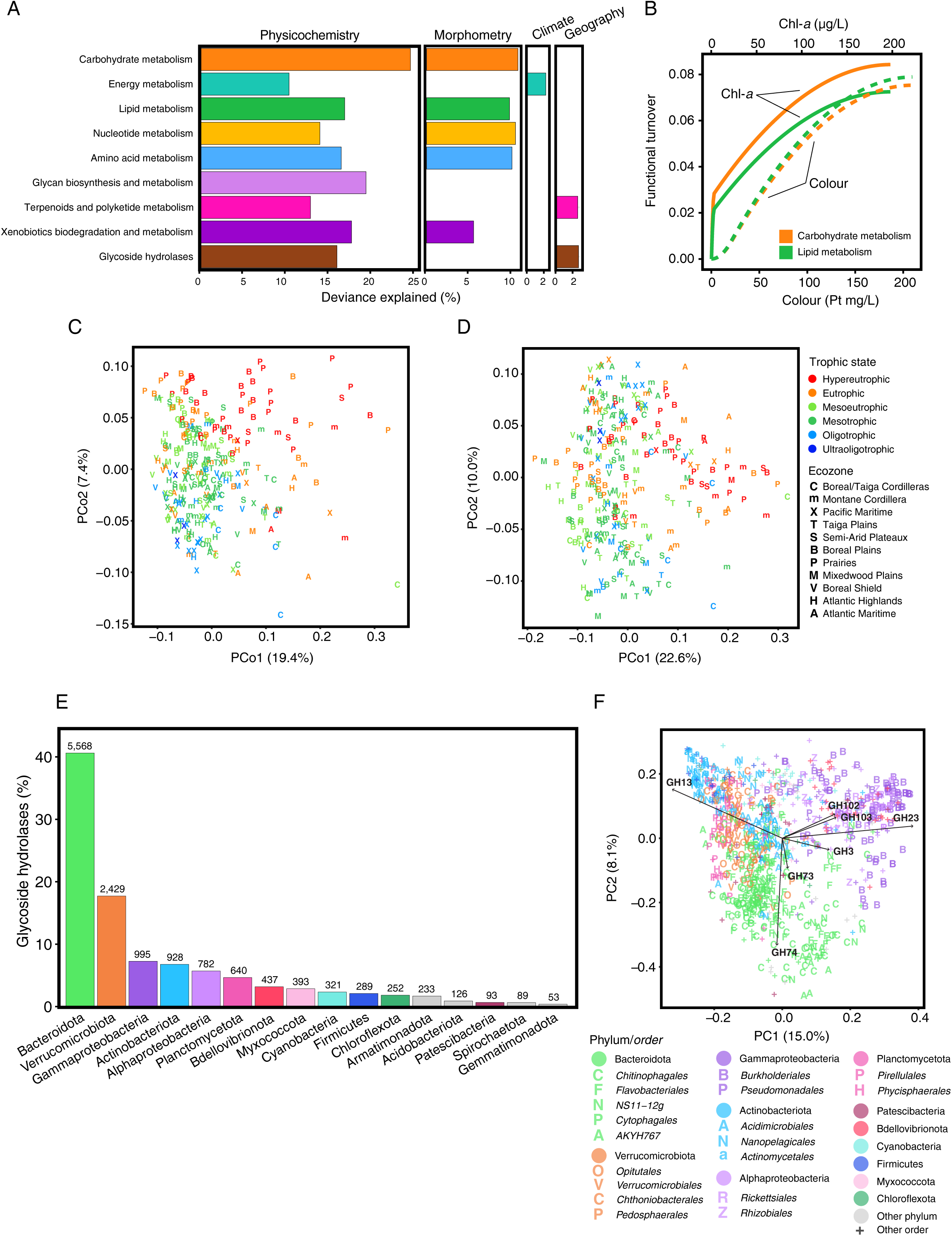
Diversity of specific bacterial functions among lake assemblages and individual MAGs. **(A)** Percent deviance explained by generalized dissimilarity models (GDMs) fitting the responses of specific MAG assemblage functions to physicochemical, morphometric, geographic, or climatic gradients. **(B)** Partial effects of chlorophyll-*a* and lake colour on the carbohydrate and lipid metabolism turnover of MAG assemblages across lakes. **(C)** PCoA showing the variation in carbohydrate metabolism among MAG assemblages. **(D)** PCoA showing the variation in lipid metabolism among MAG assemblages. Point colours in **C** and **D** represent lake trophic state and letter symbols represent ecozones. **(E)** Percentage and number of glycoside hydrolase (GH) genes distributed across MAG phyla. **(F)** PCA showing the variation in GH gene families among MAGs. Point colours represent MAG phyla and letter symbols represent order-level taxonomy within phyla.

The metabolism of complex carbohydrates of aquatic or terrestrial origin depends on initial degradation through carbohydrate-active enzymes (CAZymes). MAGs contained >13,000 genes assigned to 129 glycoside hydrolase (GH) gene families, implicated in cleaving the glycosidic bonds of diverse polysaccharides. Like carbohydrate metabolism (KEGG category), GH repertoires within bacterial assemblages were linked to lake physicochemistry (**Fig. 4A**). 41% of GH genes were identified in *Bacteroidota* MAGs (**Fig. 4E**) and were often encoded in polysaccharide utilization loci (PULs) (**Fig. S9**), demonstrating that freshwater *Bacteroidota* are central to complex organic matter degradation comparable to their role in marine and gut ecosystems^58^. *Verrucomicrobiota* MAGs were also replete with GH genes (18%), expanding on previous observations that freshwater *Verrucomicrobiota* are central players in organic matter degradation^59^. PCA of GH distributions across MAGs demonstrated a strong taxonomic structure for carbohydrate-processing potential in lake bacteria (**Fig. 4F**). *Bacteroidota* displayed the most distinct GH family repertoires, characterized by an enrichment in GH74 implicated in cellulose and xyloglucan degradation and GH20 implicated in anhydromuropeptides recycling and chitin degradation. *Proteobacteria* and *Bdellovibrionota* displayed overlapping GH repertoires distinct from other taxonomic groups and driven by GH23, GH102, and GH103, which are all lysozymes implicated in peptidoglycan degradation, aligning with potential predatory activity against other bacteria^60^. The thousands of GH genes encoded in the LakePulse MAG catalogue represent a rich resource to further examine organic matter degradation in freshwaters and to expand on previous global explorations of CAZyme diversity across ecosystems^61, 62^.

### Connecting lake bacterial diversity to watershed characteristics and human impact

While the influence of lake physicochemistry on bacterial ecology has received considerable attention and is elaborated upon in this study, terrestrial influences from the surrounding watershed are less understood. The LakePulse survey was complemented with geomatics descriptions of lake watersheds, including soils and land use^31^. Here, we coupled the geomatics data with the MAG catalogue to provide continental-scale insights into watershed impacts on lake bacterial assemblages across the land-water interface. Soil properties explained significant GDM deviance in both taxonomic (MAGs 12.1%; ASVs 23.2%) and functional (MAGs 16.3%; metagenomes 9.2%) turnover (**Fig. 5A**). Notably, across all GDMs developed in this study, the importance of soil was second only to lake physicochemistry, providing evidence for significant terrestrial influence on lake bacterial assemblage structure and function. Turnover across MAG and community profiles was most strongly associated with soil pH and the volumetric fraction of coarse fragments, a variable describing soil texture (**Fig. 5B**; **Fig. S8A**). Soil texture governs water retention, while pH determines the mobility of ions and nutrients^63^. Soil attributes thereby direct the composition and release of catchment runoff, which in turn modifies lake physicochemistry. Hence, terrestrial inputs from soils appear to influence lake bacterial diversity in a moderately predictable manner at continental scale.

**Figure 5.**
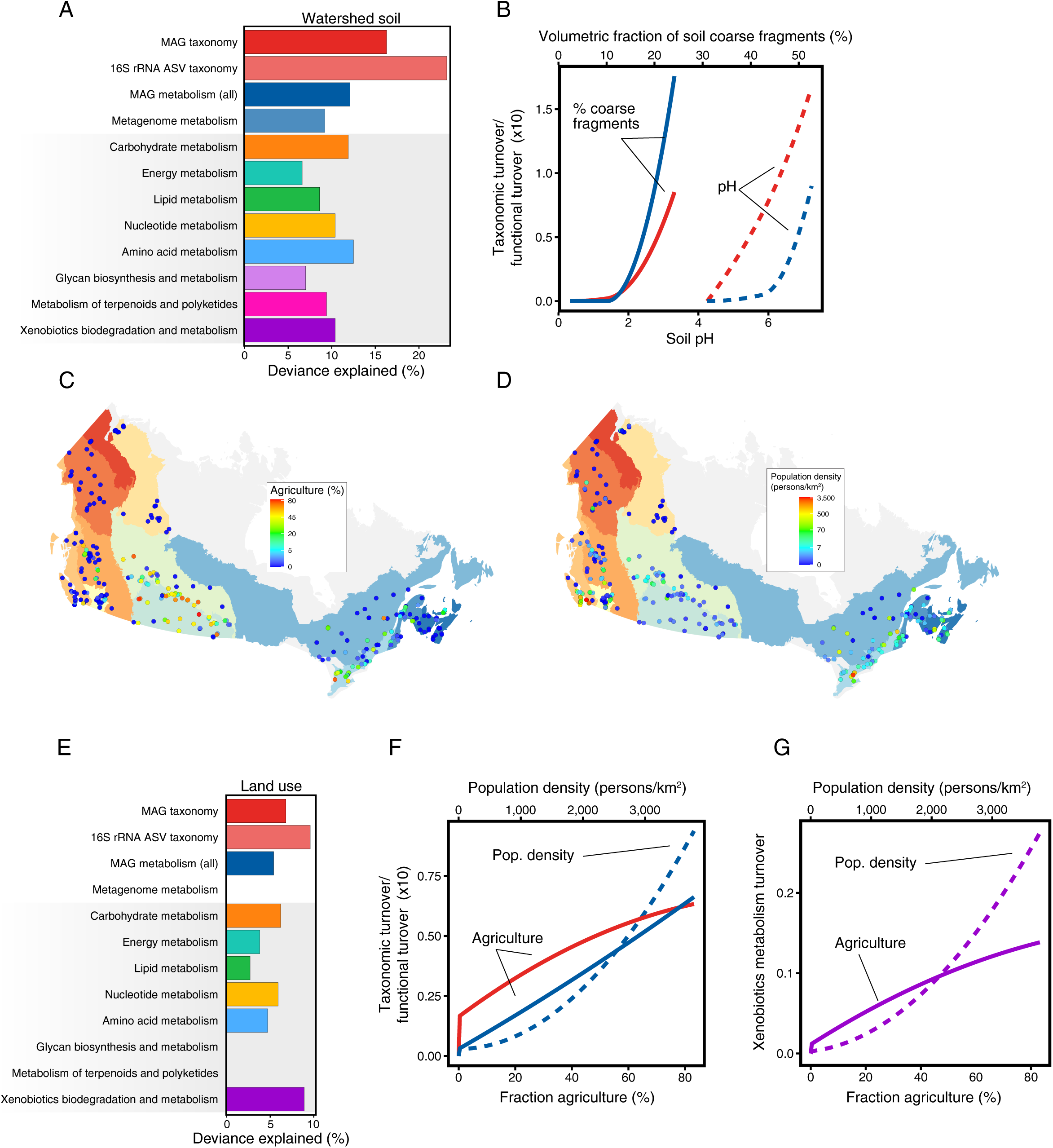
Influences of watershed soil and land use on lake bacterial assemblages. **(A)** Percent deviance explained by generalized dissimilarity models (GDMs) fitting the responses of taxonomic or functional MAG assemblages or community profiles to watershed soil characteristics. **(B)** Partial effects of soil pH and volumetric fraction of coarse fragments on the taxonomic and functional turnover of MAG assemblages across lakes. **(C)** Map of proportion of land used for crop agriculture in lake watersheds. **(D)** Map of human population densities in lake watersheds. **(E)** Percent deviance explained by generalized dissimilarity models (GDMs) fitting the responses of taxonomic or functional MAG assemblages or community profiles to watershed land use characteristics. Partial effects of crop agriculture and human population density **(F)** on the taxonomic and functional turnover of MAG assemblages across lakes and **(G)** on the turnover of xenobiotics biodegradation and metabolism within MAG assemblages across lakes.

Next, we investigated if bacterial assemblages could be predicted by human impact on the watershed. There is accumulating evidence that the taxonomic diversity of lake bacteria is influenced by land use at large spatial scales^32, 64, 65^, yet effects on function are unclear. We were particularly interested in how agriculture (**Fig. 5C**) and human population density (**Fig. 5D**) within watersheds may shape bacterial diversity, as each contribute to the eutrophication and chemical contamination of lakes. Land use GDMs explained a lesser, but still significant, amount of taxonomic (MAGs 6.8%; ASVs 9.6%) and functional (MAGs 5.4%) turnover compared to soils. There was no statistically significant response of metagenome function to land use gradients, and given that MAGs are an incomplete representation of the broader community, signals in MAG functional turnover should be interpreted cautiously. Based on evidence from GDMs, MAG assemblages are an informative subset through which to investigate – and potentially indicate – anthropogenic pressures on watersheds altering lake bacterial assemblages across the land-water interface. The functional effects of land use on MAG assemblages may reflect the responses of taxonomic composition, yet the coupling of taxonomy and function in MAGs present a dual-faceted resource to study human impact.

Of all human impact variables, agriculture exhibited a significant effect on both MAG taxonomic and functional turnover, while human population density exhibited a significant effect on functional turnover (**Fig. 5E**). Intriguingly, when analyzing specific metabolic categories, xenobiotic metabolism was the one most strongly explained by land use (**Fig. 5E**). Consistent with general functional turnover (**Fig. 5F**; **Fig. S8A**), human population density followed by agriculture were the strongest predictors of turnover within xenobiotic metabolism (**Fig. 5G**; **Fig. S8B**). In both cases, the most rapid turnover occurred where human populations were densest (1,000 – 3,000 individuals/km^2^), corresponding to the few lakes sampled within the Vancouver and Toronto metropolitan areas (**Fig. 5D**). The underlying effect of population density appeared complex: bacterial assemblages within highly populated landscapes were characterized by a lower diversity of KOs linked to xenobiotic metabolism. Moreover, PCoA showed that although xenobiotic metabolism in lakes within high-population watersheds were distinct from those within low-population watersheds, there was also large variation in xenobiotic metabolism among lakes near population centres (**Fig. S10**). As many of the detected KOs within xenobiotic metabolism were involved in the degradation of aromatic compounds, lake bacteria may be capable of metabolizing aromatic environmental pollutants, such as agricultural pesticides or industrial wastes^66^. The metabolism of land-derived pollution by lake bacteria merits further investigation, considering watersheds are increasingly converted to agriculture and urban development^67^, and altered precipitation patterns are increasing runoff^68^. Overall, these results suggest that the size and activities of human populations have a detectable and significant influence on the functional diversity encoded in the freshwater microbiome, as represented in the LakePulse MAG catalogue.

## Discussion

The LakePulse MAG catalogue, compiled from a continental-scale sampling of hundreds of lakes, addresses an urgent need to probe the microbiome underpinning freshwater ecosystems under rapid and accelerating anthropogenic change. This study has substantially expanded the number and diversity of genomes from freshwater ecosystems and demonstrated that lake bacteria are equipped to metabolize diverse substrates originating from both within lakes and their catchments. Trophic state emerged as a leading driver of taxonomic and functional variation among lake bacterial assemblages at continental scale, with implications for understanding bacterial responses to ongoing anthropogenic eutrophication of lakes^9^. The functional responses of MAG assemblages to land use pressures present a useful system for investigating human impact on the lake microbiome across the land-water interface. The links between lake bacterial diversity and watershed variables, more stable over short timescales than lake physicochemical conditions, could be leveraged to predict freshwater assemblages using remote sensing data to decrease the dependence on *in situ* surveys and monitoring^37, 40^. Our findings make clear that lake ecosystems cannot be studied in isolation of their watersheds in an era of intensifying land use^67^.

Through its construction and analysis, the LakePulse MAG catalogue makes strides toward resolving the rich and spatially heterogeneous freshwater fraction of the global microbiome. Among its potential future utilities are as an essential resource for mapping the distributions of freshwater bacteria, tracing evolutionary transitions within the land-ocean continuum, and tracking microbial dispersal and gene transfer from sources of pollution (e.g., xenobiotics; antibiotic resistance^39^) in human-altered watersheds. More lake bacterial diversity and ecosystem function remain to be captured in stratified water columns^20, 29^, sediments^69, 70^, across seasons^71^, and potentially through deeper metagenome binning^21^. Finally, microdiversity^72^ contained in the MAG catalogue may provide insights into intraspecific metabolic adaptations to the environmental heterogeneity among lakes and watersheds, including under future scenarios of environmental change.

## Methods

### Lake sampling

Lake sampling was conducted by the Natural Sciences and Engineering Research Council (NSERC) Canadian Lake Pulse Network (LakePulse) between 2017 – 2019^31^. Sampling was conducted in late June to mid-September, when some lakes are thermally stratified. To capture the natural and human-mediated heterogeneity in lake and watershed conditions, lakes were selected across three lake area categories (small: 0.1 – 0.5 km^2^, medium: 0.5 – 5 km^2^, and large: 5 – 100 km^2^) and human impact categories (low, moderate, and high as determined by land use types and coverage of the watershed) in 12 terrestrial ecozones of Canada.

Integrated surface water samples were collected over the euphotic zone up to 2 m below the surface at the site of maximum lake depth using an integrated tube sampler^73^. The euphotic zone was estimated to be twice the Secchi disk depth. Water samples were stored in chilled coolers until they were filtered onshore on the same day. Water was pre-filtered through 100 µm nylon mesh and vacuum-filtered through 0.22 µm Durapore membranes in glass funnels at a maximum pressure of 8 inHg. 500 mL of water was filtered or until the filter nearly clogged. All sampling equipment was acid-washed and rinsed with lake water prior to use. Filters were stored in sterile cryovials at −80 °C.

### Environmental data

Six categories of environmental explanatory variables were selected for analysis: (1) geography, (2) lake morphometry, (3) surface water physicochemistry, (4) watershed surface soil, (5) land use, and (6) climate. Geography variables included latitude, longitude, and altitude. Lake morphometry variables included lake area, circularity, volume, maximum depth, discharge, water residence time, watershed area, lake-to-watershed area ratio, and watershed slope within 100 m of the shoreline (data on volume, discharge, residence time, and slope were accessed from HydroLakes v. 1.0^30^). Shoreline length and average depth were strongly colinear with other morphometry variables (*r* ≥0.7) and therefore excluded from the analysis. Surface water physicochemical variables included temperature, pH, colour, and concentrations of Chl-*a*, DIC, DOC, TN, TP, calcium, chloride, magnesium, potassium, sodium, and sulfate. Missing physicochemical data were replaced with ecozone medians. Watershed soil variables estimated for the top 0 – 5 cm soil depth interval were accessed from SoilGrids250m^74^ and included bulk density of the fine earth fraction, cation exchange capacity, volumetric fraction of coarse fragments, proportions of clay, sand, and silt particles in the fine earth fraction, total nitrogen, pH, soil organic carbon content in the fine earth fraction, and organic carbon density. Land use variables were calculated as fractions of watershed area not covered by water and included crop agriculture, pasture, forestry, built development, human population density, livestock density, and poultry density. Climate variables measured over the seven days prior to lake sampling were accessed from ERA5-Land hourly reanalysis^75^ and included mean air temperature, total precipitations, mean net solar radiation, mean wind speed, and ice disappearance day for the year of sampling. Lake trophic state was categorized by TP concentrations according to the Canadian Water Quality guidelines: ultraoligotrophic (<4 μg/L), oligotrophic (4 – 10 μg/L), mesotrophic (10 – 20 μg/L), mesoeutrophic (20 – 35 μg/L), eutrophic (35 – 100 μg/L), and hypereutrophic

(>100 μg/L)^76^.

### DNA extraction and metagenome sequencing

DNA was extracted from filters using the DNeasy PowerWater kit (QIAGEN) according to the manufacturer’s instructions supplemented by the optional addition of 1 μL ribonuclease A and 30 min incubation at 37 °C. DNA was submitted to Genome Quebec for library preparation using the NEBNext Ultra II DNA Library Prep Kit (New England Biolabs) and 150 bp paired-end shotgun sequencing on an Illumina NovaSeq 6000 platform.

### Metagenome assembly and binning

Adapter-clipping and quality-trimming of raw reads were performed in Trimmomatic v. 0.36 using a Phred quality score of 33 and default settings^77^. We used a co-assembly approach to leverage the differential depth of coverage across contigs and metagenomes and thus maximize the recovery of genomes common to the same ecozone. Metagenomes were co-assembled in MEGAHIT v. 1.2.7 using k-mer lengths 27, 37, 47, 57, 67, 77, 87 and a minimum k-mer multiplicity filter of two^78^. Because only three lakes were sampled in the Taiga Cordillera and these bordered the Boreal Taiga, an ecozone with similar climate, landform, and vegetation, metagenomes from these two ecozones were combined into one co-assembly.

Local alignment of metagenome reads to co-assemblies was performed with the Burrows-Wheeler Aligner (BWA) v. 0.7.17 using the BWT-Smith-Waterman (BWTSW) index algorithm and BWA-Maximal Exact Matches^79^. Binning was performed in MetaBAT v. 2.12.1 using the default minimum scaffold length of 2,500 bp and based on the average differential coverage of each contig determined by the jgi_summarize_bam_contig_depths script implementing a 97% sequence identity threshold^80^. Bin completeness, contamination, and strain heterogeneity were estimated in CheckM v. 1.0.7^81^. Bins scoring ≥50% completeness, <10% contamination, and <10% strain heterogeneity were retained as MAGs. MAGs were dereplicated at average nucleotide identity (ANI) ≥95% and alignment coverage ≥10% in dRep v. 3.2.0 implementing gANI as the second clustering algorithm^82^. In cases where more than one MAG belonged to the same genomospecies, the highest-quality MAG was selected as the genomospecies representative to be included in the final MAG catalogue and subsequent analysis. MAGs were categorized according to MIMAG standards by which high-quality drafts display completeness >90%, contamination <5%, and ≥18/20 tRNA genes (16S, 23S, and 5S rRNA gene presence criteria were omitted) and medium-quality drafts display completeness ≥50% and contamination <10%^43^.

### Functional analysis

Protein-coding genes were annotated in Prokka v. 1.12^83^ which implemented Prodigal v. 2.6.3 for gene prediction^84^. Ribosomal RNA genes were predicted in Infernal v. 1.1.2^85^ against Rfam v. 14.2^86^. Gene functions were annotated in KofamScan using default settings and a bitscore-to-threshold ratio of 0.7^87^. Transporters were predicted in BLASTp against the Transporter Classification Database downloaded on January 27, 2021^88^ using a maximum e-value of 10^-20^. CAZymes were predicted in hmmsearch from HMMER v. 3.1b2^89^ against dbCAN2 HMMdb v. 9^90^. GH family 109 was omitted from CAZyme analysis due to nonspecificity of the hidden Markov model.

Functions were assigned based on the following set of rules. Oxygenic photosynthesis required evidence of at least one marker gene from both photosystems I (psaA-F: K02689, K02690, K02691, K02692, K02693, or K02694) and II (psbA-F: K02703, K02704, K02705, K02706, K02707, or K02708). For anoxygenic photosynthesis, evidence of photosystem II required the presence of both pufL (K08928) and pufM (K08929); evidence of photosystem I required the presence of all four proteins (pscA-D: K08940, K08941, K08942, K08943). Bacteriorhodopsin assignment required evidence of bop (K04641). Autotrophic carbon fixation via three independent pathways was evidenced by rbcL (K01601) and prkB (K00855) for the CBB cycle, mct (K14470) for the glyoxylate assimilation pathway, and abfD (K14534) for the 3HP/4HB cycle. Evidence of glycan import mediated by the susCD protein complex was determined by the presence of both susC (K21573) and susD (K21572). Aromatic compound degradation was determined by the presence of either a funneling or ring fission pathway: evidence of catabolic funneling was identified by ferulate to vanillin conversion (evidenced by fcs (K12508)), vanillin to protocatechuate conversion (evidenced by vanA (K03862)), or protocatechuate ring opening (evidenced by pcaG (K00448) or ligA (K04100)); ring fission was identified by gallate ring opening (evidenced by galA (K04099)), catechol ring opening (evidenced by K00446 or K03381), salicylate to gentisate conversion (evidenced by K00480 or K18242 or K18243), or gentisate ring opening (evidenced by K00450). Methanol oxidation was evidenced by xoxF (K23995). Nitrogen fixation was evidenced by nifD (K02586) or nifH (K02588) or nifK (K02591). Dissimilatory nitrate reduction was evidenced by narG (K00370) and narH (K00371) or napA (K02567) and napB (K02568). Evidence of nitrite reduction was determined by the presence of 1) nirB (K00362) and nirD (K00363), 2) nrfA (K03385) and nrfH (K15876), 3) nirK (K00368), or 4) nirS (K15864). Nitric oxide reduction was evidenced by norB (K04561) and norC (K02305). Nitrous oxide reduction was evidenced by nosZ (K00376). Evidence of sulfur oxidation via SOX was determined by the presence of 1) soxA (K17222) and soxB (K17224) and soxX (K17223) and soxY (K17226) and soxZ (K17227), or 2) soxC (K17225) and soxD (K22622). Dissimilatory sulfate reduction was evidenced by 1) aprA (K00394) and aprB (K00395), or 2) dsrA (K11180) and dsrB (K11181).

PULs were delineated following the algorithm described in Terrapon *et al*.^91^. First, adjacent SusC and SusD genes located on the same DNA strand (termed tandem susCD pairs) were identified. Tandem susCD pairs within five loci of each other were considered a single susCD-containing locus. Next, operons structured around tandem susCD pairs were delineated by including genes within intergenic distances of ≤200 bp. PUL boundaries were iteratively extended to neighbouring operons when a CAZyme was detected within the first five genes. The selected intergenic distance threshold exceeded the empirically-derived 102 bp cut-off proposed by Terrapon *et al*.^91^ so as to include functionally-associated genes other than CAZymes that would otherwise be excluded. Only CAZymes implicated in glycan degradation (glycoside hydrolases, polysaccharide lyases, carbohydrate-binding modules, and carbohydrate esterases) were included in conditional PUL extension. Finally, only PULs containing specified CAZyme types were retained.

### MAG read recruitment and coverage normalization

To avoid recruitment to conserved regions, rRNA and tRNA gene sequences were masked in BEDTools v. 2.26.0^92^ prior to mapping. To determine the coverage of MAGs across all lake metagenomes, trimmed metagenome reads were initially recruited to MAGs at a sequence identity threshold of 95% in BBMap v. 37.76^93^, then filtered with a hard cut-off of 96% identity using a custom Python script. The selection of a 95% alignment similarity cut-off reflected a prokaryotic genome ANI species delineation^94^. Mapping files were converted from SAM to sorted BAM format in Samtools^95^. Horizontal coverage was calculated following the method of Rodriguez-R *et al*.^21^ as TAD_80_, the average sequencing depth truncated to the central 80% of mapped positions normalized by the number of genome equivalents. Genome equivalents in each metagenome were estimated in MicrobeCensus v. 1.1.0^96^. A positive TAD_80_ value signified that ≥10% of MAG positions were mapped by metagenome reads; the ≥10% sequencing breadth threshold has been shown to detect species presence at a confidence of >95%^97^.

### Taxonomic classification and phylogenetic analysis

MAG taxonomic classification was performed in GTDB-Tk v. 1.3.0 by phylogenetic placement into bacterial reference tree v. r95^98^. A *de novo* phylogenetic tree was constructed in GTDB-Tk using the alignment of 120 marker genes based on the JTT model of evolution and subsequently plotted with the R package ggtree^99^.

### Metagenome analysis

Single metagenome assemblies were generated in MEGAHIT^78^ using k-mer lengths 23, 43, 63, 83, 103, 123 or metaSPAdes^100^ v. 3.13.0 using k-mer lengths 21, 33, 55, 77, 99, 127 and mapped in BWA v. 0.7.17 using the BWTSW index algorithm and BWA-Maximal Exact Matches^79^. Assemblies and coverage files were submitted to the DOE-JGI Metagenome Annotation Pipeline v. 4.16.6, 5.0.20, or 5.0.23^101^. Metagenome functional diversity was quantified by relative gene counts, calculated as KO counts normalized by total contig average depth of coverage.

### 16S rRNA gene amplicon sequencing and analysis

A ∼300 bp fragment of the V4 region of the 16S rRNA gene was amplified using the primer set 515F (5’-GTGCCAGCMGCCGCGGTAA-3’) and 806R (5’-GGACTACHVGGGTWTCTAAT-3’)^102^. Each PCR mixture consisted of 5 µL Phusion High Fidelity Buffer, 0.5 µL dNTPs (10 mM), 1.8 µL of each primer (5 µM), 0.25 µL Phusion polymerase, 13.65 µL H_2_0, and 10 ng in a 2 µL volume of template DNA. PCR conditions were 30 s of 98 °C, followed by 22 cycles of 98 °C for 20 s, 54 °C for 35 s, 72 °C for 30 s, and a final elongation at 72 °C for 60 s. Sequencing of 250 bp paired-end reads was performed on an Illumina MiSeq platform.

Primer sequences were trimmed in Cutadapt v. 3.1^103^. ASVs were inferred in DADA2 v. 1.16^104^, which performed trimming of low-quality 3’-end positions from forward and reverse reads, quality retention of reads 160 – 200 bp in length, ASV inference within individual samples informed by machine-learned sequencing error rates, merging of paired reads, and removal of consensus chimeric ASVs. ASVs <250 or >256 bp in length were removed along with ASVs not assigned at the kingdom level and ASVs assigned to archaea, eukaryotes, or chloroplasts. ASV data from two lakes were discarded because they did not meet the threshold 10,000 sequences. ASVs were classified against GTDB r95 16S rRNA gene sequences formatted for DADA2^105^.

### Ecological analysis

PCA of lake features was performed in the R package vegan v. 2.5.6^106^ on scaled and centred variables; latitude was passively plotted. Biogeographical analysis was conducted on MAG assemblages in 300 freshwater and oligosaline lakes, identified as having conductivity <8 mS/cm and total major ions <4,000 mg/L. To build MAG functional assemblage matrices, we assumed that a function was implied when an encoding MAG was mapped to a lake metagenome. PCoA was performed in the R package ape v. 5.4^107^ on pairwise Jaccard dissimilarities between MAG or ASV taxonomic assemblage presence-absence data or Bray-Curtis dissimilarities between metagenome gene count data normalized by total depth of coverage. GDMs fitting assemblage turnover to environmental gradients were constructed in the R package gdm v. 1.4.2 using 100 permutations during each step of backward variable elimination^108^. PCA of MAG GH composition was performed on Hellinger-transformed data. Data analysis and visualization were performed in R v. 4.0.1^109^ using the tidyverse v. 1.3.0 suite of packages^110^.

### Code availability

Scripts associated with this study are available at https://github.com/rebeccagarner/lakepulse_mags.

### Data availability

Raw metagenome reads were archived in the European Nucleotide Archive under study accession PRJEB29238 (https://www.ebi.ac.uk/ena/browser/view/PRJEB29238). Metagenome co-assemblies were deposited and annotated at the Joint Genome Institute (JGI) Genomes OnLine Database (GOLD) under study accession <u>Gs0136026</u> and analysis projects Ga0495746 (Boreal/Taiga Cordilleras), Ga0495744 (Montane Cordillera), Ga0495745 (Pacific Maritime), Ga0495743 (Taiga Plains), Ga0485099 (Semi-Arid Plateaux), Ga0485102 (Boreal Plains), Ga0485100 (Prairies), Ga0364548 (Mixedwood Plains), Ga0373103 (Boreal Shield), Ga0372599 (Atlantic Highlands), and Ga0372598 (Atlantic Maritime). MAGs from co-assemblies and annotations were deposited in Dryad (https://doi.org/10.5061/dryad.zkh1893fs).

## Acknowledgements

This study was funded by the NSERC Canadian LakePulse Network (Strategic Network Grant NETGP-479720) and Canada Research Chairs held by DAW and YH. REG and VEO received scholarships from the NSERC CREATE ÉcoLac Training Program in Lake and Fluvial Ecology. REG also received support from a *Fonds de recherche du Québec – Nature et technologies* Doctoral Research Scholarship and the Stephen Bronfman Graduate Scholarship in Environmental Studies. The pan-Canadian field campaigns were made possible through the immense commitment and efforts of the LakePulse sampling crews and support teams, and the cooperation and assistance of Indigenous groups, municipal and park employees, lake associations, and landowners. We thank the LakePulse researchers involved in the generation, management, and quality control of the dataset, and acknowledge contributions from Jelena Juric, Paul MacKeigan, Cindy Paquette, Marieke Beaulieu, Katherine Griffiths, Geneviève Potvin, Bruno Cremella, Gabriel Diab, Vincent Fugère, Jihyeon Kim, Anaïs Oliva, and Katherine Velghe. We thank Wentworth Brookes, Shawn Simpson, and Compute Canada for bioinformatics technical support.

## Supplementary information

**Figure S1.** Project workflow illustrating the generation and analysis of the LakePulse MAG catalogue.

**Figure S2.** Distributions of **(A)** geography, **(B)** lake morphometry, **(C)** watershed soil, **(D)** land use, **(E)** climate, and **(F)** surface water physicochemistry variables. Colours represent ecozones and ecozone medians are indicated by dashed lines.

**Figure S3.** Number of MAGs generated within each ecozone co-assembly.

**Figure S4.** Comparison of taxonomic diversity in MAG and 16S rRNA gene ASV datasets: percentages of MAGs and ASVs assigned to **(A)** phyla and **(B)** orders.

**Figure S5.** Phylum-level biogeographic distributions of MAGs.

**Figure S6.** Hierarchical clustering (Ward’s linkage) of MAG phylum-level taxonomic assemblages. Letter symbols above each MAG assemblage (scaled to relative TAD_80_) represent ecozones and coloured squares represent lake chlorophyll-*a* concentration.

**Figure S7.** PCoAs of **(A)** the taxonomic variation among ASV assemblages based on Jaccard dissimilarities and **(B)** the variation in metabolic gene content among metagenomes based on Bray-Curtis dissimilarities.

**Figure S8.** Significant predictors of **(A)** taxonomic and functional turnover across MAG assemblages and community profiles and **(B)** turnover in specific metabolic functions across MAG assemblages. The height of bars represents the relative importance of predictors within the generalized dissimilarity model. MAG assemblage turnover is shown above zero and community (ASV or metagenome) turnover is shown below zero.

**Figure S9.** Gene maps of polysaccharide utilization loci (PULs) identified across *Bacteroidota* MAGs. Arrows indicate gene directions. Colours represent gene types (SusCD, CAZymes, tRNA genes, other KOs).

**Figure S10.** Principal coordinate analysis (PCoA) showing the variation in xenobiotics biodegradation and metabolism among MAG assemblages.

**Table S1.** Summary of metagenome information: lake names, accession information, sampling coordinates, and assembly characteristics.

**Table S2.** Summary of MAG accession information, quality, genome characteristics, taxonomy, associated file names, marker gene content, and TAD_80_ across 300 freshwater to oligosaline lakes.

**Table S3**. Number of novel MAGs in each phylum.

**Table S4.** Generalized dissimilarity modelling results for MAG assemblages and community (ASV and metagenome) profiles across lakes based on taxonomic and functional composition.

