## Supplementary figures and images for "A genome catalogue of lake bacteria across watershed land use gradients at a continental scale"

### Figure S1

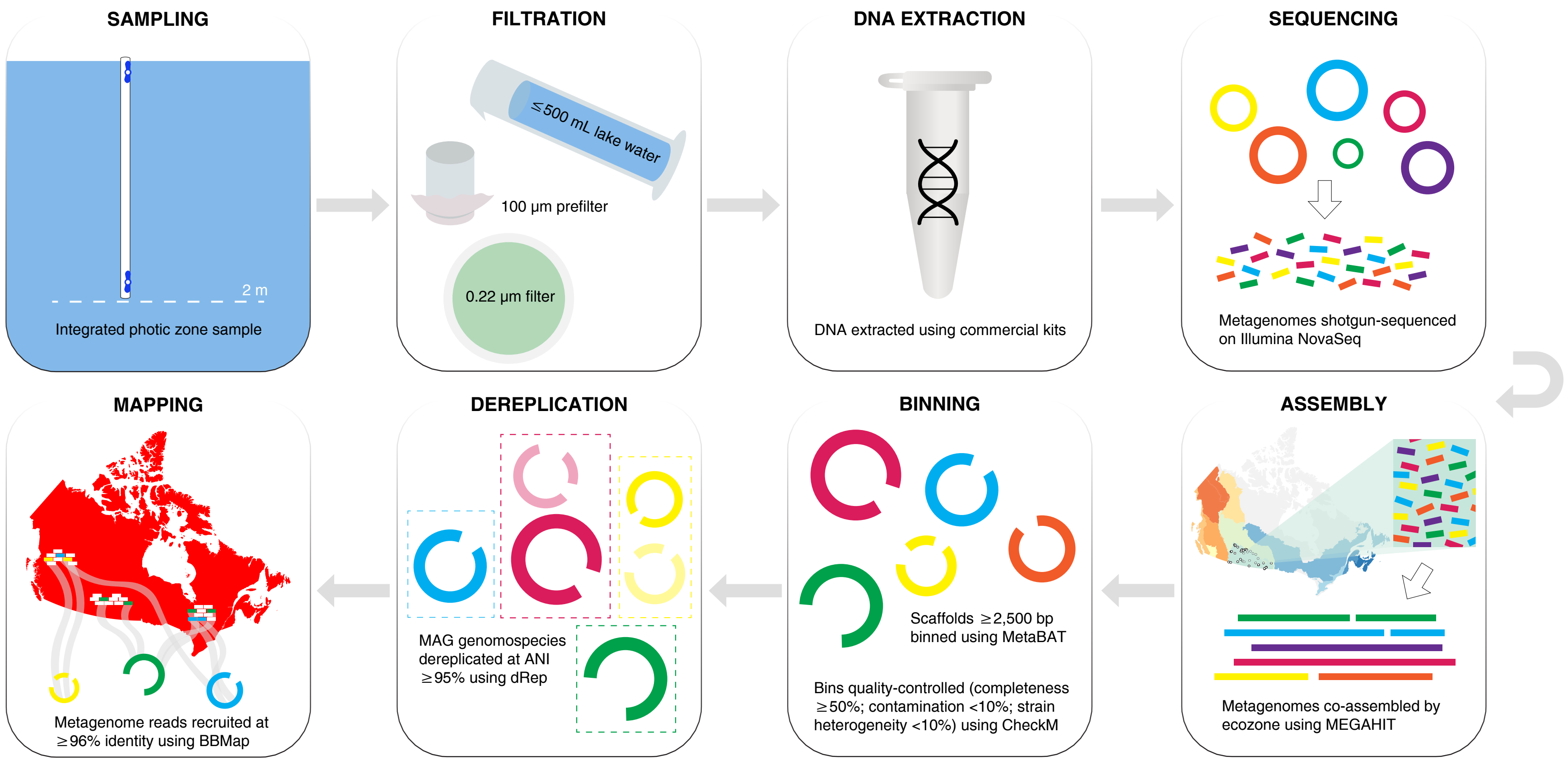

### Figure S2

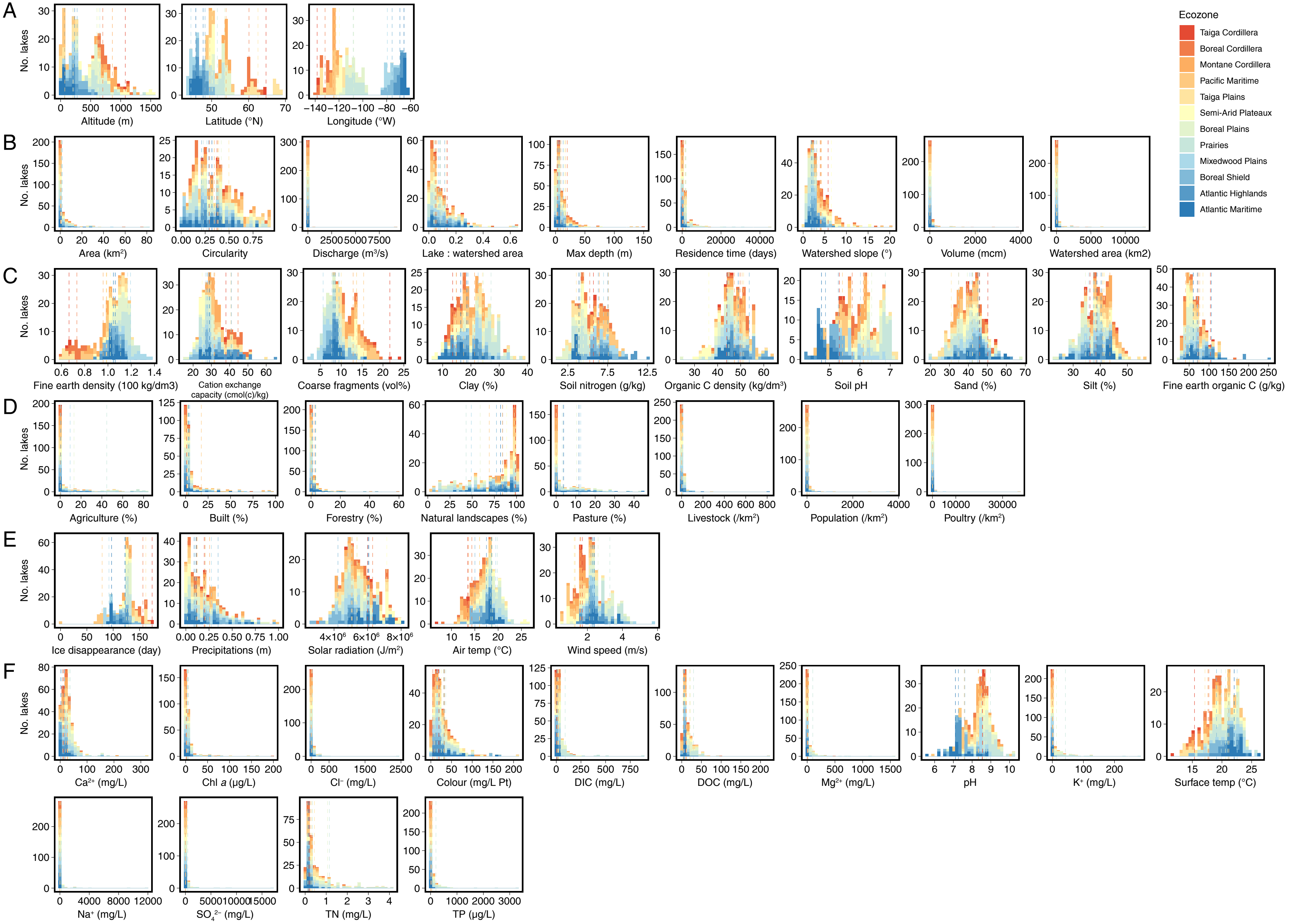

### Figure S3

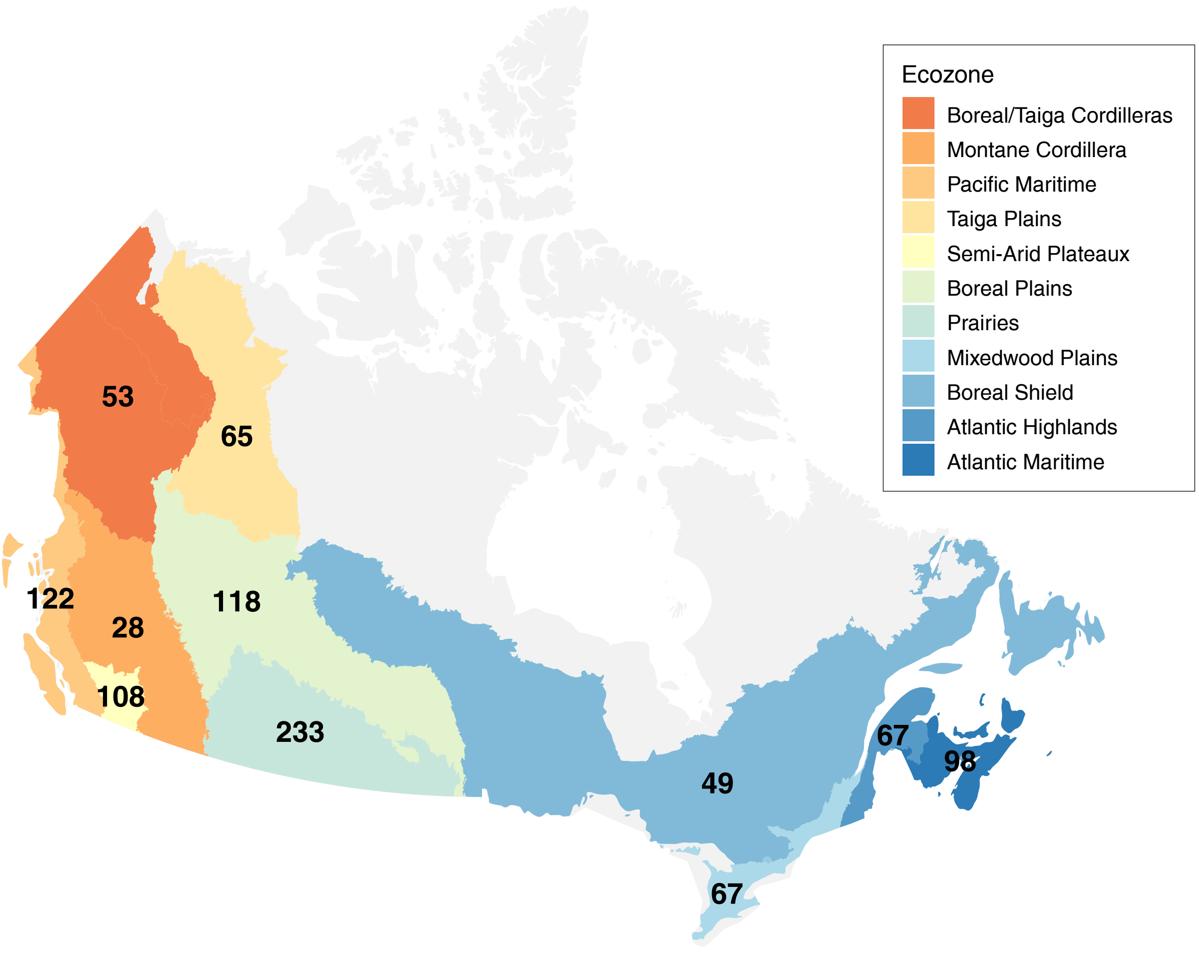

### Figure S4

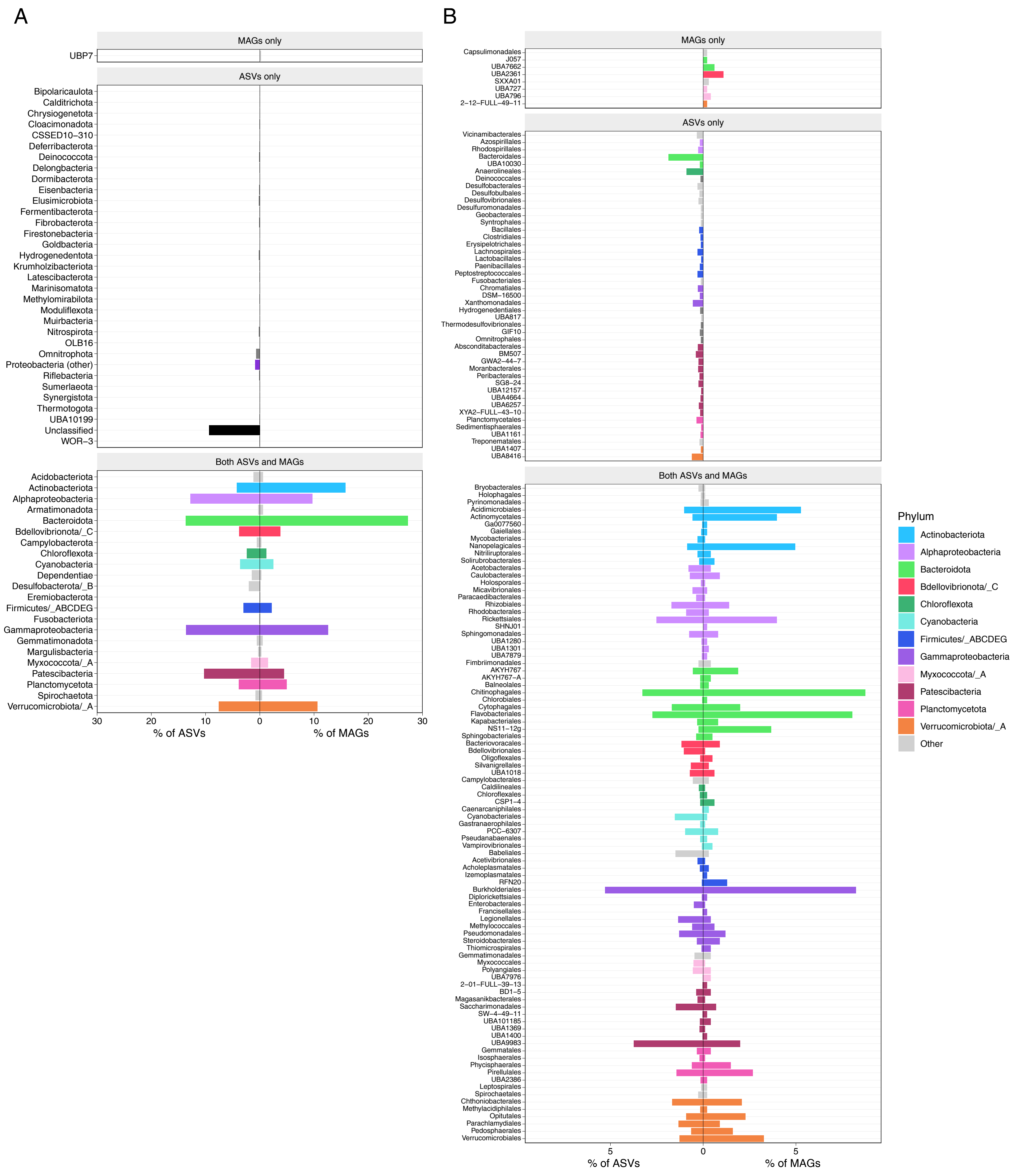

### Figure S5

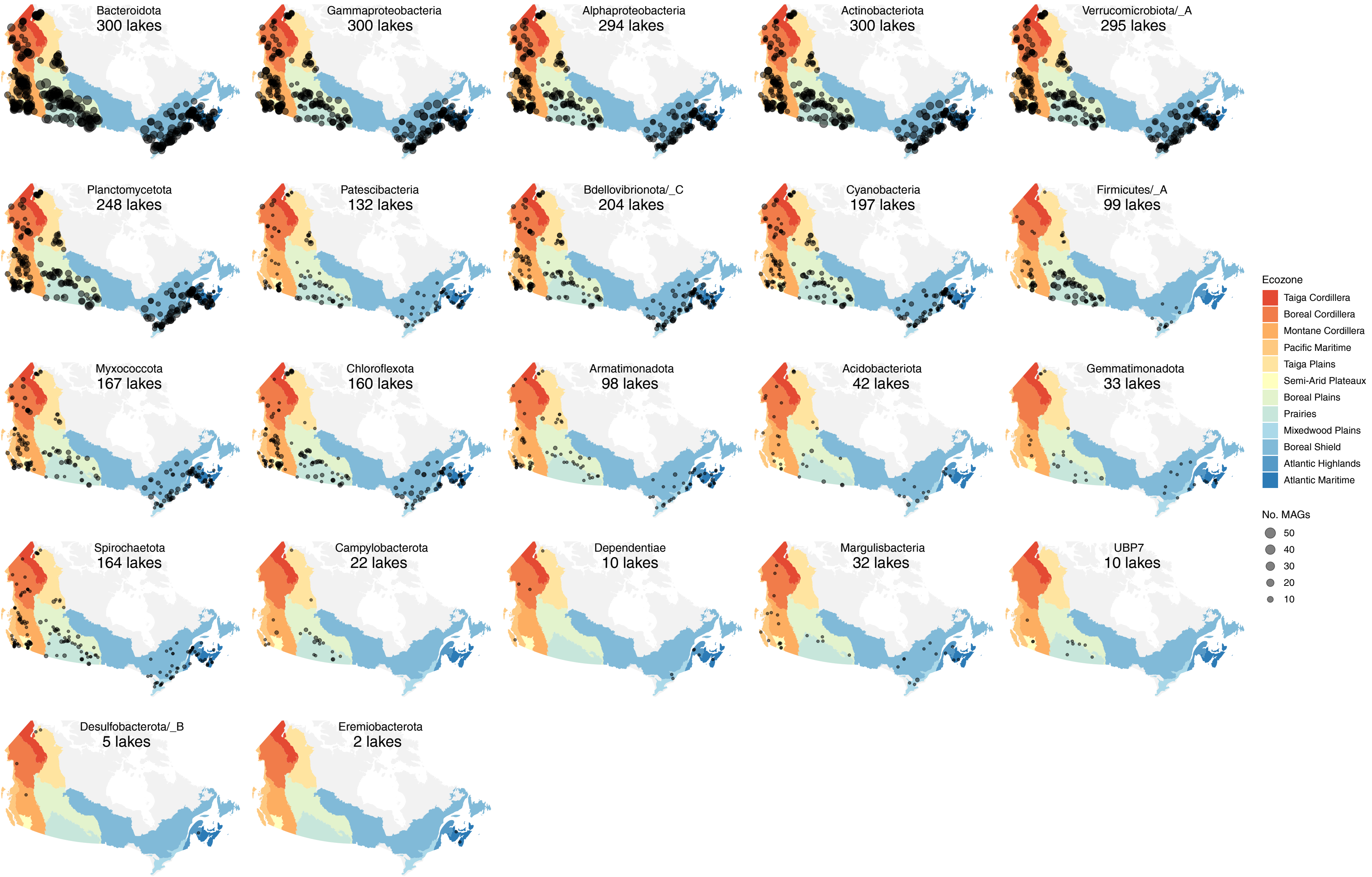

### Figure S6

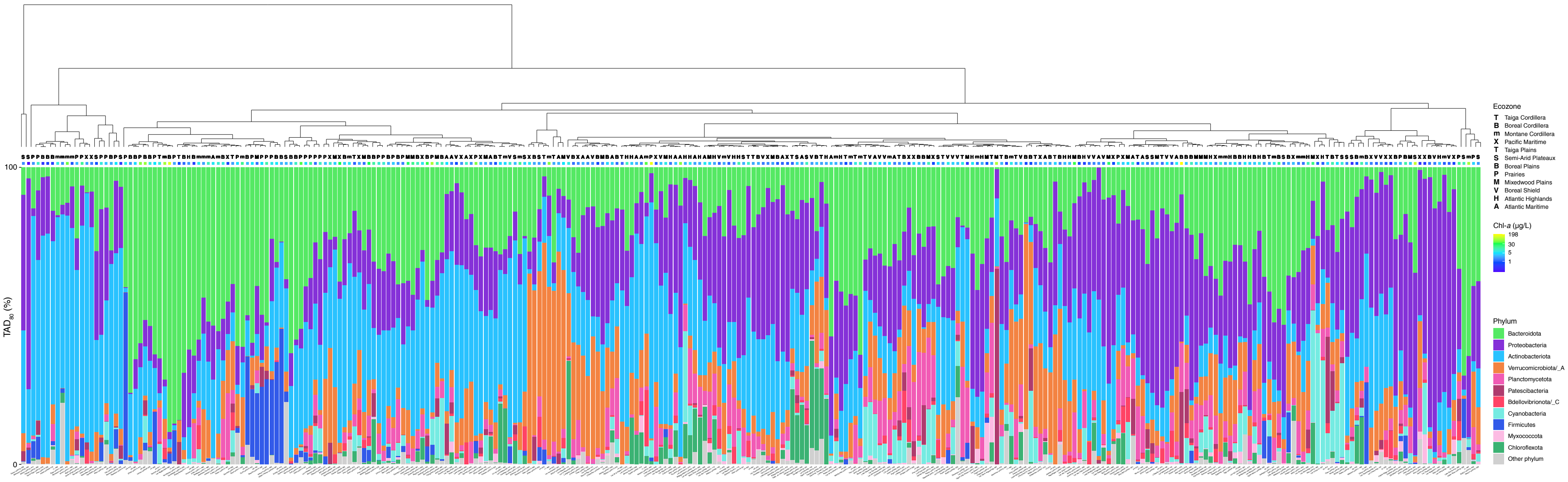

### Figure S7

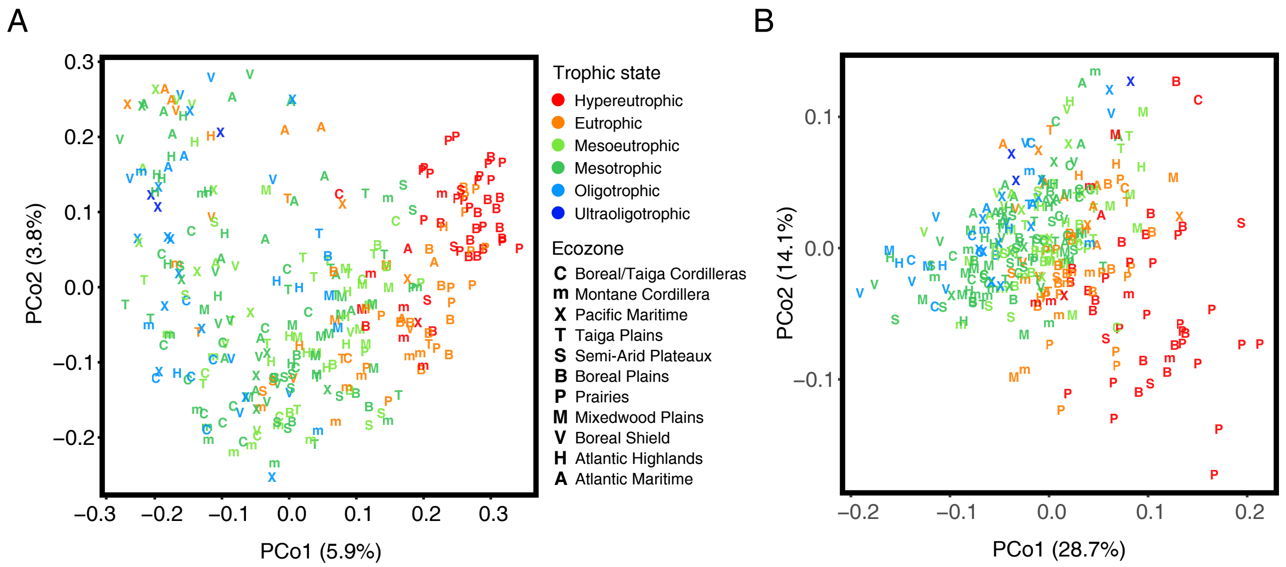

### Figure S8

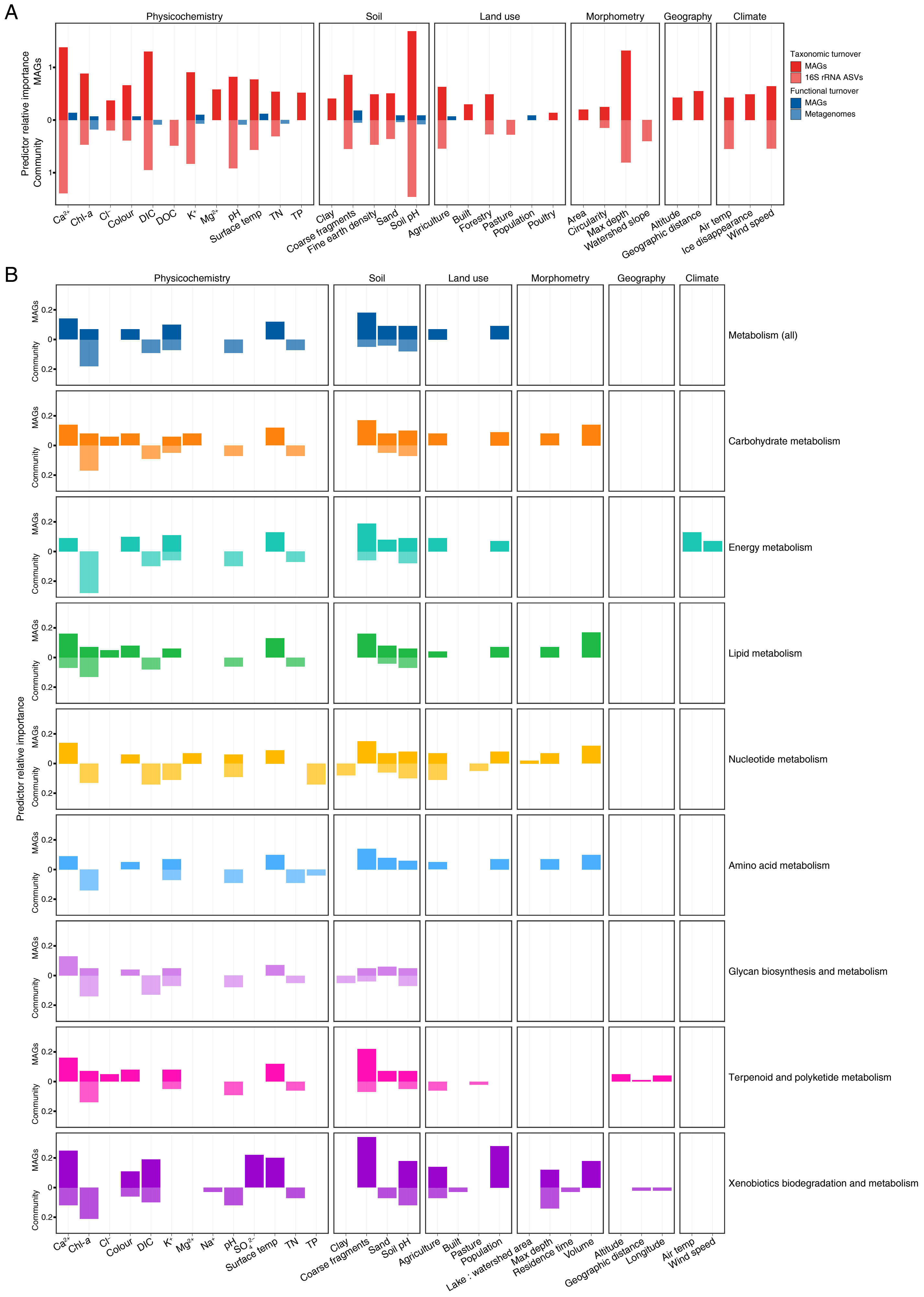

### Figure S9

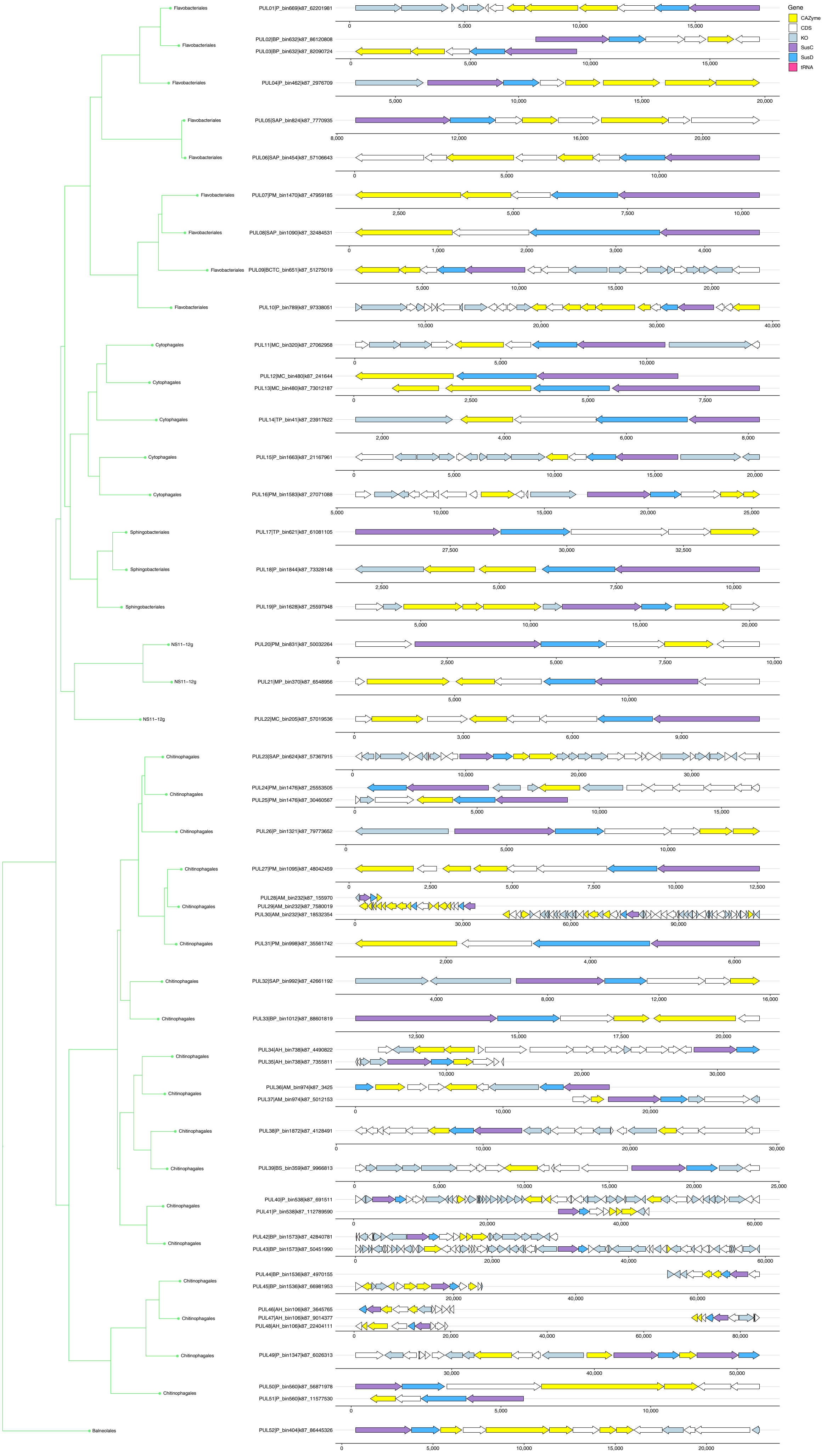

### Figure S10

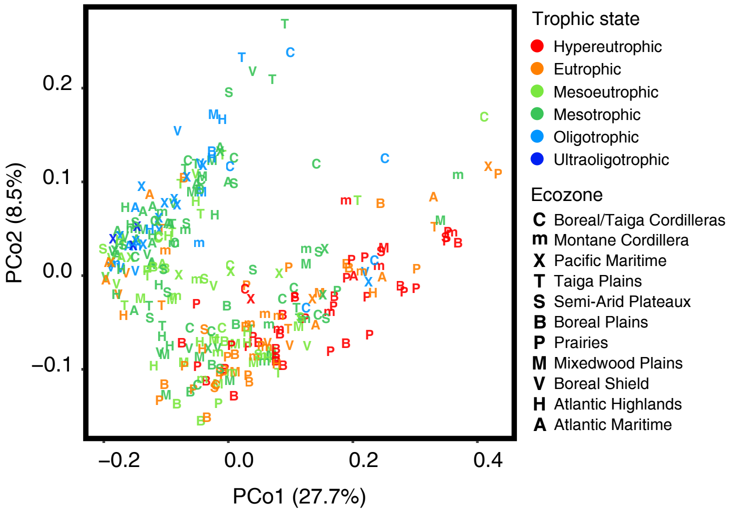
